# Characterizing the body morphology of the first metacarpal in the Homininae using 3D geometric morphometrics

**DOI:** 10.1101/2020.04.30.070326

**Authors:** Jonathan Morley, Ana Bucchi, Carlos Lorenzo, Thomas A. Püschel

## Abstract

**Objectives:** The morphological characteristics of the thumb are of particular interest due to its fundamental role in enhanced manipulation. Despite its possible importance regarding this issue, the body of the first metacarcapal (MC1) has not been fully characterized using morphometrics. This could provide further insights into its anatomy, as well as its relationship with manipulative capabilities. Hence, this study quantifies the shape of the MC1’s body in the extant Homininae and some fossil hominins to provide a better characterization of its morphology.

**Materials and methods:** The sample includes MC1s of modern humans (n=42), gorillas (n=27) and chimpanzees (n=30), as well as *Homo neanderthalensis, Homo naledi* and *Australopithecus sediba*. 3D geometric morphometrics were used to quantify the shape of MC1’s body.

**Results:** The results show a clear distinction among the three extant genera. *H. neanderthalensis* mostly falls within the modern human range of variation. *H. naledi* varies slightly from modern humans, although also showing some unique trait combination, whereas *A. sediba* varies to an even greater extent. When classified using a discriminant analysis, the three fossils are categorized within the *Homo* group. Conclusion: The modern human MC1 is characterized by a distinct suite of traits, not present to the same extent in the great apes, that are consistent with an ability to use forceful precision grip. This morphology was also found to align very closely with that of *H. neanderthalensis. H. naledi* shows a number of human-like adaptations, whilst *A. sediba* presents a mix of both derived and more primitive traits.

## 1 Introduction

Thumbs in modern humans are different from those of African apes (e.g., Almécija, Smaers, & Jungers, 2015; Dunmore, Bardo, Skinner, & Kivell, 2020; Galletta, Stephens, Bardo, Kivell, & Marchi, 2019; Green & Gordon, 2008; Stephens et al., 2016; Susman, 1994). Modern humans have a relatively broader shaft for the first metacarpal (MC1) and a higher thumb-to-digit ratio than the African apes, especially chimpanzees (Almécija et al., 2015; Feix, Kivell, Pouydebat, & Dollar, 2015; Green & Gordon, 2008; Rolian & Gordon, 2013). Additionally, compared to our closest living relatives we possess thenar musculature that is relatively more developed than the other hand muscles (Tuttle, 1969), a condition that has been inferred from the hominin fossil record by observing how strong or flawed their bony attachments are (Bush, Lovejoy, Johanson, & Coppens, 1982; Karakostis, Hotz, Tourloukis, & Harvati, 2018; Kivell, 2015; Kivell, Kibii, Churchill, Schmid, & Berger, 2011; Maki & Trinkaus, 2011; Richmond et al., 2020). Modern humans also differ from the extant African apes in the relative size of the epicondyles and degree of curvature of the proximal (Marchi, Proctor, Huston, Nicholas, & Fischer, 2017; Marzke et al., 2010) and distal joint surfaces of the MC1 (Galletta et al., 2019).

These anatomical traits that set apart humans from the African apes have presumably evolved to cope with the different functional demands experienced by these taxa (i.e., manipulation vs. locomotion) (Almécija, Moyà-Solà, & Alba, 2010; Matarazzo, 2015; Püschel, Marcé-Nogué, Chamberlain, Yoxall, & Sellers, 2020; Richmond & Strait, 2000; Tsegai et al., 2013). The more robust human thumb and greater degree of curvature of the joint surfaces allow our species to produce greater force and to better withstand the stresses of tool-related behaviors (Galletta et al., 2019; Key & Dunmore, 2018; Key & Dunmore, 2015; Rolian, Lieberman, & Zermeno, 2011). On the other hand, the thumb of chimpanzees is slender and shorter relative to the other fingers, presumably a suspensory-related adaptation (Almécija et al., 2015; Feix et al., 2015), and although the thumb length and breadth in gorillas differs less from humans than chimpanzees (Almécija et al., 2015; Feix et al., 2015; Green & Gordon, 2008), it does show a reduced thenar musculature, which is the primitive condition in the hominidae (Diogo, Richmond, & Wood, 2012; Tocheri, Orr, Jacofsky, & Marzke, 2008; Tuttle, 1969).

Even though using skeletal proxies of the MC1 to infer the degree of dexterity is common practice (e.g., Dunmore et al., 2020; Feix et al., 2015; Maki & Trinkaus, 2011), the continuous nature of these traits makes it difficult to quantify how different hominines are with respect to each other, which consequently complicates the correlation of these proxies with different functional capabilities. Building upon this problem, recent research on the MC1 has been conducted using three-dimensional geometric morphometric (3DGM) techniques, focusing on the joint surfaces in apes and fossil hominins (Galletta et al., 2019; Marchi et al., 2017). Marchi et al. (2017) propose that hominins (modern humans, *Paranthropus robustus/early Homo* SK84 and *Au. africanus*) are significantly different from non-human hominids in that they possess a radioulnar and dorsovolar flatter proximal joint, a less projecting volar beak and a radially extended surface. This would allow our species to better abduct and to accommodate larger axial loads when pinching objects (Marchi et al., 2017; Marzke et al., 2010). Humans also vary from apes in having a larger and flatter distal articular surface in a radioulnar direction and a radial palmar condyle that is larger and more palmarly projecting than the ulnar one, which would contribute to the stabilization of the joint during forceful precision grip (Galletta et al., 2019). Neanderthals and *H. naledi* are located within the modern human range of variation for these traits, whereas the other analyzed hominins (*Au. africanus, Paranthropus robustus/early Homo* SK84 and *Au. sediba*) occupy a position between modern humans and the great apes.

In spite of its possible importance, the body of the MC1 has not been fully analyzed using 3DGM to assess its possible relevance when correlating its anatomy with different manipulative capabilities. In addition, fossils are often fragmentary and epiphyses in the fossil record are often damaged (see for e.g., *H. naled*’s U.W. 101-401 MC1) or abraded (see for e.g., *H. naled*’s U.W. 101-1641 MC1). Therefore, a method focused only on the MC1 shaft might be particularly useful. Consequently, this study focuses on the body morphology of the MC1 using 3DGM. The objective was to provide further information that could contribute towards a better characterization of the MC1’s anatomy, as well as to provide further insights towards the identification of structures in extant species that may be associated with human-like manipulative capabilities and to assess if similar morphologies are present in fossil hominins.

Even though the great apes use their hand for manipulatory activities, their morphology is likely more related to their locomotion (i.e., knuckle-walking and arborealism) (Almécija, Moyà-Solà, & Alba, 2010; Matarazzo, 2015; Püschel et al., 2020; Richmond & Strait, 2000; Tsegai et al., 2013). It is therefore expected that the selective pressures associated with locomotor behavior in chimpanzees and gorillas will result in an MC1 morphology that varies significantly from that of modern humans. We also expect gorillas to be closer to humans rather than chimpanzees, as previous research has indicated that their metacarpals are broader and the thumb-fingers ratio less different from humans compared with those of chimpanzees (Almécija et al., 2015; Green & Gordon, 2008; Rolian & Gordon, 2013). Additionally, we also expect the MC1’s shaft morphology of *Au. sediba* to show an intermediate morphology located between the range of variation of modern humans and that of the great apes, as previous studies indicate that the hand of this species displays a mosaic anatomy of primitive and derived traits (Kivell et al., 2011). *Au. sediba* MC1 has gracile attachments for the opponens pollicis and first dorsal interosseus muscles, but it also possesses a long thumb relative to the fingers, which is close to the modern human configuration (Kivell et al., 2011). On the other hand, we expect that the Neanderthal and *H. naledi* specimens will show a morphology similar to that of modern humans as previous analyses have suggested that they exhibit similar attachment sites of the thenar musculature, as well as a relatively similar thumb length (Feix et al., 2015; Karakostis, Hotz, Tourloukis, & Harvati, 2018; Kivell, 2015; Maki & Trinkaus, 2011). Consequently, we tested the following hypothesis:

### Hypothesis 1

MC1’s morphology significantly differs between modern humans and the extant African ape species. The modern human MC1’s shaft is expected to be more similar to that of gorillas rather than chimpanzees due to its broader shaft, as well as relative length and breadth.

### Hypothesis 2

*H. naledi* and *H. neanderthalensis* specimens exhibit an MC1 morphology more similar to modern humans than other great apes, while *Au. sediba* shows an intermediate morphology between the African apes and modern humans.

## 2 Material and methods

### 2.1 Sample

The extant sample used in this study includes MC1s of modern humans (*Homo sapiens*; n=42), chimpanzees (*Pan troglodytes*; n=30), and gorillas (*Gorilla gorilla* and *Gorilla beringei*; n=27) (Table S1). The human MC1s came from a medieval cemetery in Burgos, Spain (Casillas Garcia & Alvarez, 2005) and the surface models were obtained using a Breuckmann SmartSCAN structured light scanner. The non-human sample came from museum collections and are of different origins (i.e., wild-shot, captivity and unknown origin). There were no significant shape differences between wild vs. captive specimens, nor between the two gorilla species included in this study, hence we felt confident to pool the extant specimens at the genus level (see S2 for further details). The MC1 surface models were collected using photogrammetry as described in Bucchi et al. (2020a). The 3D models from the surface scanner were obtained using a resolution of 0.125 mm, while most of the photogrammetric models ranged from 400,000 to 600,000 triangles of uniform size. A previous study that applied the same surface scan and photogrammetry protocols to digitize hand bones found that both types of 3D models are of comparable quality (Bucchi, Luengo, Bove, & Lorenzo, 2020b). Additionally, we carried out a comparison a sub-sample of 30 specimens that were digitized using both technologies (i.e., photogrammetry and structured-light scanning) and we found that differences between models obtained using the different digitalization technologies are extremely small (less than ~0.17 mm on average). Hence, we are confident that it is possible to combine these kinds of 3D models in our analyses. Further details about these 3D model comparisons can be found in S3. Only adult individuals that show no evident pathologies were included in the study and right MC1s were preferred (although left MC1s were reflected when their antimere was not present as indicated in S1).

The fossil sample includes the right metacarpal from a *Homo neanderthalensis*, the right metacarpal from a *Homo naledi* and the left metacarpal from an *Australopithecus sediba*. The *H. neanderthalensis* sample (La Ferrassie 1) was found in La Ferrassie archaeological site in Savignac-de-Miremont, France. This skeleton was discovered in 1909 and is estimated to be 70–50,000 years old (Guérin et al., 2015). The *Homo naledi* sample (U.W. 101-1321) was recovered in 2013 from the Rising Star cave system in South Africa and has been dated to around 236-335 ka years ago (Dirks et al., 2017). The *A. sediba* sample (MH2) was taken from the near complete wrist and hand of an adult female (Malapa Hominin 2) discovered in Malapa, South Africa, which has been dated around 1.98 million years (Berger et al., 2010; Pickering et al., 2011). The latter fossils were downloaded from Morphosource https://www.morphosource.org/, whereas the Neanderthal was obtained from a cast housed at the Catalan Institute of Human Paleoecology and Social Evolution (IPHES).

### 2.2. 3DGM

3D landmarks were collected using the software Landmark Editor 3.6 (Wiley et al., 2005) to quantify the MC1’s morphology, including relevant functional proxies as the epicondyles, the shaft curvature and the attachments sites for the opponens pollicis, abductor pollicis longus and first dorsal interosseus muscles. These attachments sites are in the MC1 at the lateral margin, body at the ulnar side of the bone and the base at the radial side, respectively, and are the same for the three genera under study (Diogo et al., 2011; 2013). Eight curves comprising 20 equidistant landmarks each were placed at pre-defined points on the MC1 (Figure 1; S4). These landmarks were chosen to provide a good representation of the shaft of the bone.

**Figure 1.**
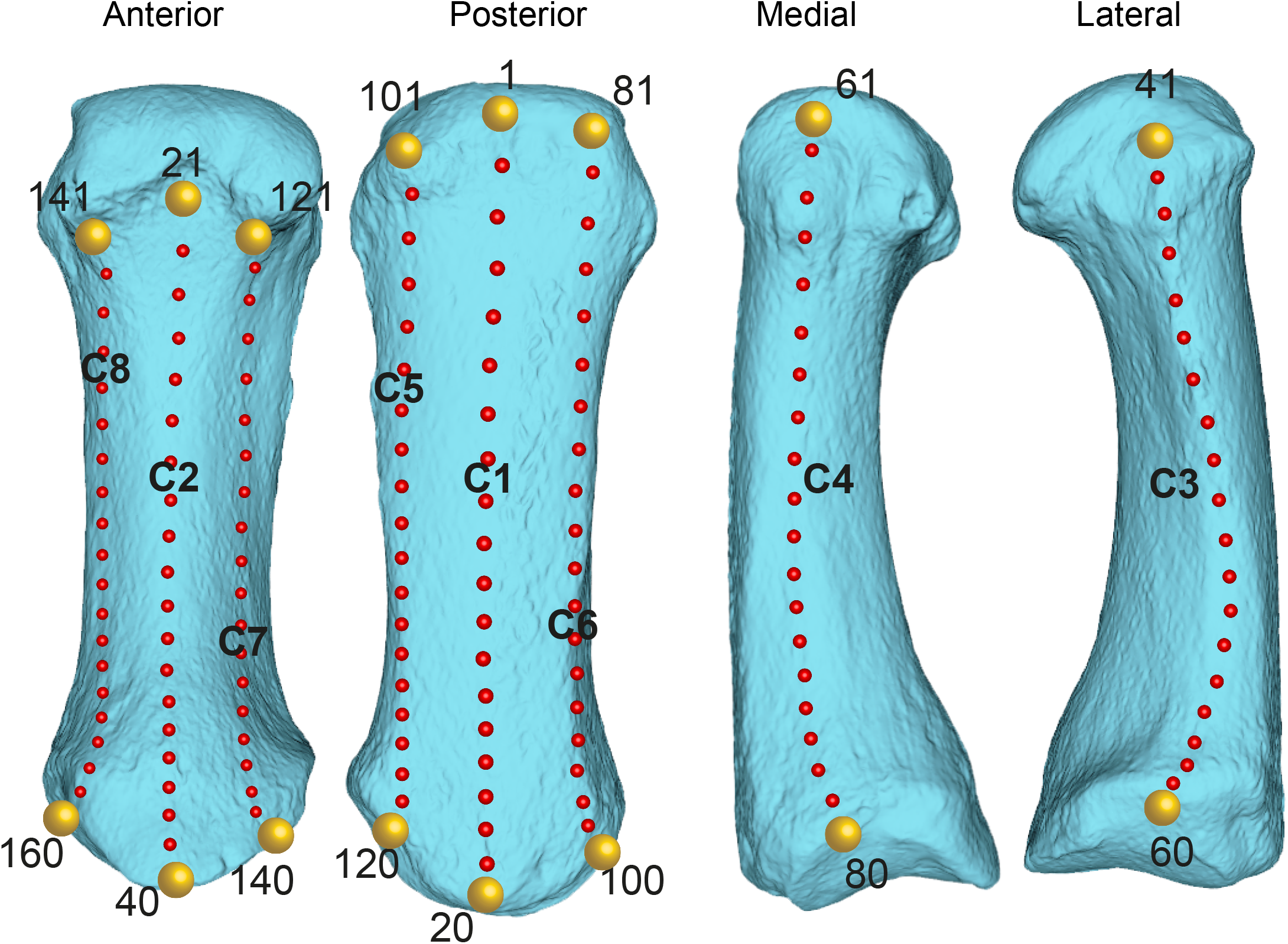
Illustration of the 16 landmarks (yellow spheres) and 144 semi-landmarks (red spheres) used in this study. The numbers of the 16 fixed landmarks and eight semi-landmark curves (C1-C8) are also plotted.

We assessed whether sufficient number of landmarks have been sampled to characterize MC1’s shape variation by using the lasec() function of the ‘LaMBDA’ 0.1.0.9000 R package (Watanabe 2017) (further details about this procedure can be found in S5). The first and last landmarks from each one of the eight curves were treated as fixed (i.e., 16 fixed landmarks), whereas all the rest of them (i.e., 144 landmarks) were considered as semi-landmarks. A generalized Procrustes superimposition was performed on the landmark data to remove differences due to scale, translation, and rotation in order to obtain shape variables (Bookstein, 1991). This procedure was done using the gpagen() function available as part of the ‘geomorph’ R package 3.3.1 (Adams, Collyer, & Kaliontzopoulou, 2020). The semi-landmarks were slid on the MC1’s surface by minimizing bending energy (Bookstein, 1997; Gunz, Mitteroecker, & Bookstein, 2005). This is an iterative process that works by allowing the semi-landmarks to slide along the surface to remove the effects of arbitrary spacing by optimizing the location of the semi-landmarks with respect to the consensus shape configuration (Gunz & Mitteroecker, 2013). There are two main criteria to slide semi-landmarks (i.e., bending energy and Procrustes distance) which have been shown to provide relatively similar results when carrying out inter-specific comparisons (Perez, Bernal, & Gonzalez, 2006). We preferred to use bending energy as this sliding criterion allows all semi-landmarks to slide together and being influenced by the other available landmarks and semi-landmarks, whereas when Procrustes distance is used, each semi-landmark slides individually and, apart from the common Procrustes superimposition (Gunz & Mitteroecker, 2013). All the Procrustes residuals analyzed in this work are available in S6.

These obtained shape variables were then used in a principal component analysis (PCA) to summarize shape variation. The PCA was carried out using the gm.prcomp() function of the ‘geomorph’ R package 3.3.1 (Adams, Collyer, & Kaliontzopoulou, 2020). To visualize shape differences warped models representing the shape changes along the first three principal components (PCs) were generated. The models closest to the mean shape (i.e., lowest Procrustes distance to the multivariate consensus) was warped to match the multivariate mean using the thin plate spline method. Then, the obtained average model was warped to display the variation along the three plotted PC axes (mag = 1).

The dataset of extant hominoids was then grouped by genus and the Procrustes variance of observations in each group (i.e., the mean squared Procrustes distance of each specimen from the mean shape of the respective group) was computed as a simple measure to assess morphological disparity within each one (Drake & Klingenberg, 2010; Zelditch, Swiderski, & Sheets, 2012a). Procrustes variance was applied here as way to evaluate intra-genus variation, and absolute differences in Procrustes variances were computed to test differences in morphological disparity among groups (these differences statistically evaluated through permutation [999 rounds]). This procedure was carried out using the morphol.disparity() function available as part of the ‘geomorph’ R package 3.3.1 (Adams, Collyer, & Kaliontzopoulou, 2020).

A multi-group linear discriminant analysis (LDA) (also known as canonical variate analysis [CVA]) was run to test if it was possible to distinguish among the three genera. This procedure maximizes the separation between groups. Since our number of variables (i.e., landmarks and semi-landmarks) exceeded the number of analyzed specimens, we carried out this analysis using the principal components (PCs) that accounted for 90% of the sample variance to reduce the dimensionality of the dataset. The LDA was carried out using the lda() function of the ‘MASS’ 7.3-51.6 R package (Venables & Ripley, 2002). Performance was calculated using the confusion matrix from which the overall classification accuracy was computed, as well as the Cohen’s Kappa statistic (Püschel, Marcé-Nogué, Gladman, et al., 2020; Püschel, Marcé-Nogué, Gladman, Bobe, & Sellers, 2018). The complete dataset was resampled using a ‘leave-one-subject-out’ cross-validation, as a way to asses classification performance (Kuhn & Johnson, 2013). In addition, by using the obtained discriminant function we classified the fossil sample into the three extant genera as way to assess morphological affinities. Pairwise PERMANOVA tests with Bonferroni corrections for multiple comparisons were performed to assess shape differences among the three extant genera using the PCs that accounted for 90% of the sample variance. Euclidean distances computed using the PCs that accounted for ~ 90% of the total variance of the sample were selected as dissimilarity index. This procedure was performed using the adonis() function of the ‘vegan’ 2.5-7 R package (Oksanen et al., 2020).

Additionally, we also decided to compute a curvature metric to better assess how curve the MC1’s shaft is along both its dorsal and palmar aspects (i.e., semi-landmark curves C1 and C2 in Fig. 1), as well as to facilitate the morphological description of the morphometric results. Hence, Menger (1930) curvatures were calculated for each one of the semi-landmark points of the two curves (i.e., C1 for the dorsal side and C2 for the palmar aspect) using a custom-written script in R. The Menger curvature of three points in n-dimensional Euclidean space 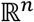 is defined as the reciprocal of the radius of the circle that passes through the three points (Menger, 1930). Menger curvature was calculated locally for each semi-landmark point along the curve, excepting the first and last fixed landmarks, as it is not possible to compute a curvature value at the starting and ending points of each one of the curves. This resulted in 18 curvature values for each one of the semi-landmark curves (i.e., C1 and C2). The curvature values of each one of the curves were summed to obtain a measurement of the overall curvature of C1 and C2 (higher values would correspond to more pronounced curvatures). This procedure was performed on the six 3D models that were warped to represent the variation along the first three PC axes.

In addition, as measurement error (ME) has a critical importance when performing morphometric analyses, a sub-sample of 33 randomly selected MC1s were digitized twice and compared via a Procrustes ANOVA to assess ME (Klingenberg & McIntyre, 1998). We also carried out a regression of shape variables on centroid size using the whole sample to assess allometric influence. Both procedures were carried out using the procD.lm() function available as part of the ‘geomorph’ R package 3.3.1 (Adams, Collyer, & Kaliontzopoulou, 2020). All the mentioned morphometric and statistical analyses were carried out in R 4.0.2 (R Core Team, 2020).

## 3 Results

### 3.1 Measurement error and allometric influence

The Procustes ANOVA used to measure intra-observer error in the sub-sample showed that the mean square for individual variation far exceeded ME, so this type of error was negligible (see S7 for further details). ME was also quantified as shape repeatability using a ratio of the among-individual to the sum of the among-individual and measurement error components as explained in Zelditch, Swiderski, & Sheets (2012b). Shape repeatability was 0.95, which indicates a minimal ~5% error. Regarding allometric influence, we found that centroid size only accounted for ~2.7% of MC1 shape variation. This means that, for the goals of the present study, we can exclude size as a particularly significant factor contributing to potential inter-generic variation in shape. Hence, we decided that it was not necessary to ‘correct’ for allometric effects as ~97.3% of the shape variation is not explained by size (further details about this regression are available in S8).

### 3.2 Principal component analysis

The PCA performed using the shape variables returned 102 PCs. The first 22 PCs accounted for ~ 90% of the total variance of the sample, hence offering a reasonable estimate of the total amount of MC1’s shape variation, which were then used in the LDA and pairwise PERMANOVA tests. The first three PCs in the PCA account for ~ 58% of the total variance and display a relatively clear separation among the extant African ape genera (Fig. 2a) (PCA biplots for PC1 vs. PC2, PC1 vs. PC3 and PC2 vs. PC3 are also available in S9). PC1 explains 41.44%, PC2 11.18% and PC3 5.82% of total variance, respectively (Fig. 2). To visualize shape differences, the warped 3D models corresponding to the highest and lowest values at each extreme of the first three PCs were plotted alongside the violin plots. A violin plot is a combination of a boxplot and a kernel density plot that is rotated and placed on each side to show the distribution shape of the data (Adler & Kelly, 2020). In addition, six movies showing the shape changes along the three first PCs axes are also provided in S10. These warped models are also displayed in Figure 3 to facilitate the morphological interpretation of our results. Anatomical descriptions associated with each one of the positive and negative extremes of the first three PCs are also provided in the same figure.

**Figure 2.**
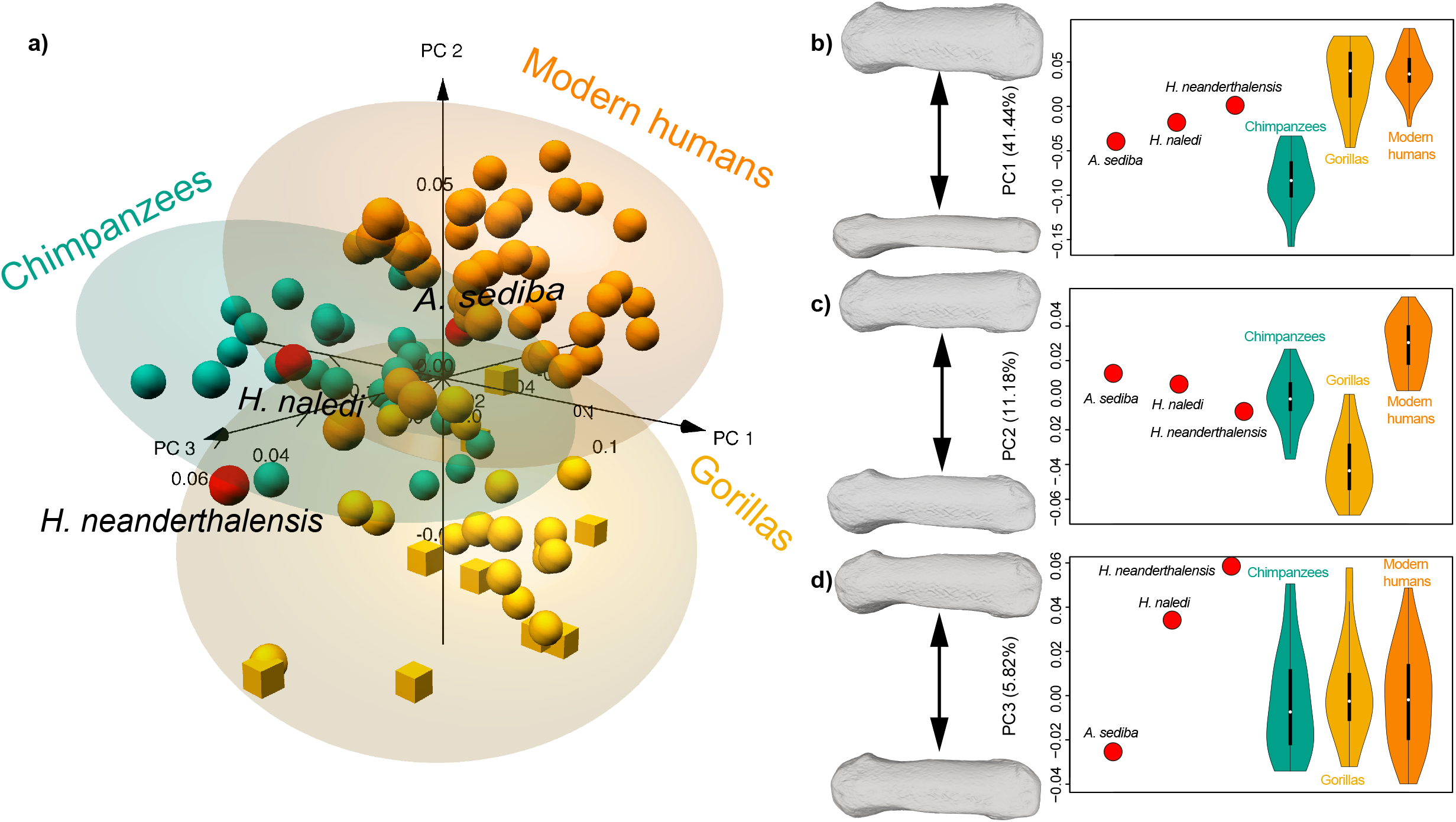
Principal component analysis of the shape data: the a) three main axes of morphological variation are displayed (ellipses represent 95% confidence intervals, red spheres: fossils, orange spheres: *H. sapiens*, green spheres: *P. troglodytes*; golden spheres: *G. gorilla*, golden cubes: *G. beringei*); Violin plots of the PCs scores of the analyzed sample are shown for b) PC1, c) PC2 and d) PC3 (fossil values are displayed as red triangles). The white dot in the middle is the median value, whilst the thick black bar in the center represents the interquartile range. The thin black line extended from it corresponds to the upper (maximum) and lower (minimum) adjacent values in the data. The distribution shape of the data for each one of the three PCs is represented by a kernel density plots that were rotated and placed on each side of each one of the boxplots. To visualize shape differences warped models representing the shape changes along the first three principal components (PCs) were plotted alongside the violin plots (dorsal views). The models closest to the mean shape was to match the multivariate mean using the thin plate spline method. Then, the obtained average model was warped to display the variation along the three plotted PC axes (mag = 1).

**Figure 3.**
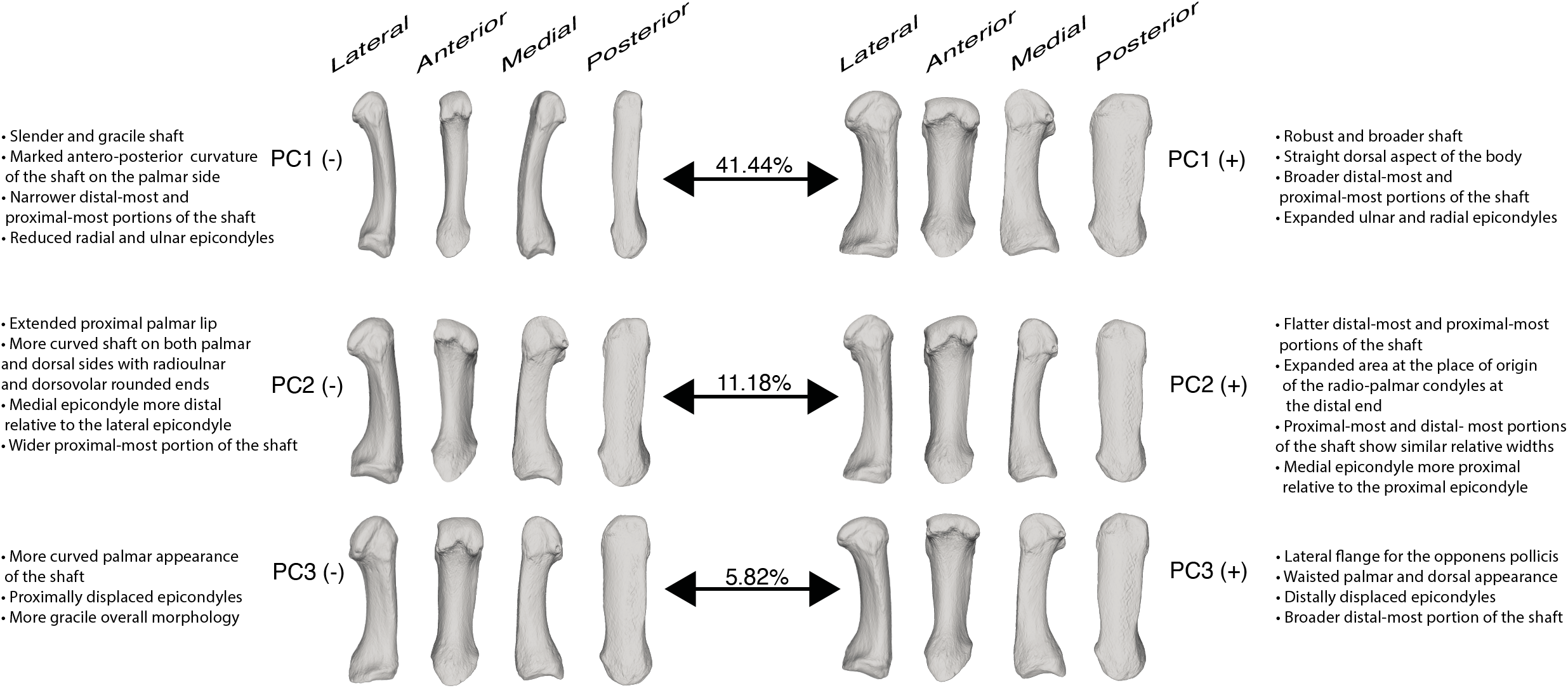
Warped models representing the shape changes along the first three principal components (PCs). The models closest to the mean shape was to match the multivariate mean using the thin plate spline method. Then, the obtained average model was warped to display the variation along the three plotted PC axes (mag = 1). Corresponding anatomical descriptions are provided alongside each one the warped models. Please notice that articular surfaces were not morphometrically characterized and as such, none of the anatomical descriptions refer to them. References to curvature on this figure are based on the results provided in Table 5.

Violin plots of PC1 (Fig. 2b) show a notable difference between gorillas and humans vs. chimpanzees. PC1 separates the mediolaterally narrower MC1 shafts of *Pan* from the broad MC1 shafts of *H. sapiens* and *Gorilla*. These two genera exhibit the highest PC1 scores, which correspond to a more developed muscular attachments, straighter dorsal aspect of the body, and overall robust shaft with broader distal-most and proximal-most portions of the shaft (i.e., MC1’s body and not the articular surfaces which were not morphometrically characterized in this study) (Fig. 3). Chimpanzees show the lowest PC1 scores, representing a more gracile shaft, a more pronounced antero-posterior curvature of the shaft, less marked muscle attachments with narrower distal-most and proximal-most portions of the body and smaller radial and ulnar epicondyles (Fig. 3). *H. neanderthalensis* falls within the human and gorilla distributions and is distinct completely from the chimpanzees. *H naledi* falls within the gorilla distribution, whilst *A. sediba* is characterized by a lower PC1 score and aligns closer to the *Pan* distribution. None of the analyzed fossils fall within any of the interquartile ranges (IQR) (i.e., black bars in Fig. 2b-d) of any of the extant genera.

Violin plots of PC2 (Fig. 2c) shows distinct variation among the extant genera, with a morphological continuum ranging from *Gorilla* (lower PC2 values), *Pan* (central PC2 values) and extant *Homo* (higher PC2 scores). PC2 seems to summarize the relative breadth of the middle and distal shaft with respect to the relative size of the proximal shaft and base. The *Gorilla* sample has the lowest PC2 scores, a more radioulnar and dorsovolar rounded ends of the shaft, and a medial epicondyle which is more distal relative to the lateral epicondyle (Fig.3). The modern human distribution shows the highest PC2 scores, representing flatter distal-most and proximal-most portions of the body, as well as larger area at the place of origin of the radial palmar condyles at the distal end of the analyzed area (i.e., the shaft). The chimpanzee sample lies in between the gorilla and modern human samples displaying an intermediate morphology. In a similar fashion as chimpanzees, the three fossils are located at intermediate positions in PC2 distribution. *H. neanderthalensis* and *H. naledi* display PC2 scores that are within the *Pan* IQR, whilst *A. sediba* has higher values.

Violin plots of PC3 (Fig. 2d) show a similar distribution of PC scores for the three extant genera. From a morphological perspective, lower values correspond to more gracile MC1s while higher scores are associated with more robust morphologies displaying more surface for muscular attachments (for the opponens pollicis, first dorsal interosseous and abductor pollicis longus muscles). *H. naledi* and *A. sediba* show values which are within the extant genera distributions, but outside their IQR and at opposite extremes of the axis. *H. neanderthalensis* lies outside the distribution of any of the extant genera, probably due to its particularly robust morphology and associated marked muscular insertion areas, in particular a marked lateral flange for the opponens pollicis.

### 3.3 Morphological disparity

The obtained results show that three extant genera show a similar magnitude of disparity. Nevertheless, gorillas exhibit a higher Procrustes variance as compared to modern humans and chimpanzees (Table 1a). Gorillas are significantly different to modern humans, and chimpanzees when comparing absolute variance differences, whilst modern human do not significantly differ from chimpanzees (Table 1b).

### 3.4 Linear discriminant analysis

The LDA model using the first 22 PCs clearly distinguishes among the three extant genera, displaying an outstanding performance with excellent classification results after cross-validation (Accuracy: 0.97; Cohen’s Kappa: 0.95; Fig. 4; Table 2). When using the obtained discriminant function to classify the fossils into the extant categories (as a way of assessing morphological affinities) (Table 3), the three of them were classified into the *Homo* category, even though only *H. neanderthalensis* was located within the 95% confidence interval of the modern humans (Fig. 4). The posterior probabilities were extremely close to 1 for *H. naledi and H. neanderthalensis*, hence indicating that, in spite of their differences, their morphology is closer to that of modern humans. *A. sediba* was also classified within the *Homo* category (posterior probability: 63%) but this specimen also showed non-trivial posterior probabilities classifying it within the *Gorilla* category (posterior probability: 29%) or as a member of the *Pan* group (posterior probability: 7%). There were significant differences among all extant genera when analyzing the 22 PCs from the PCA carried out using the shape variables (Table 4).

**Figure 4.**
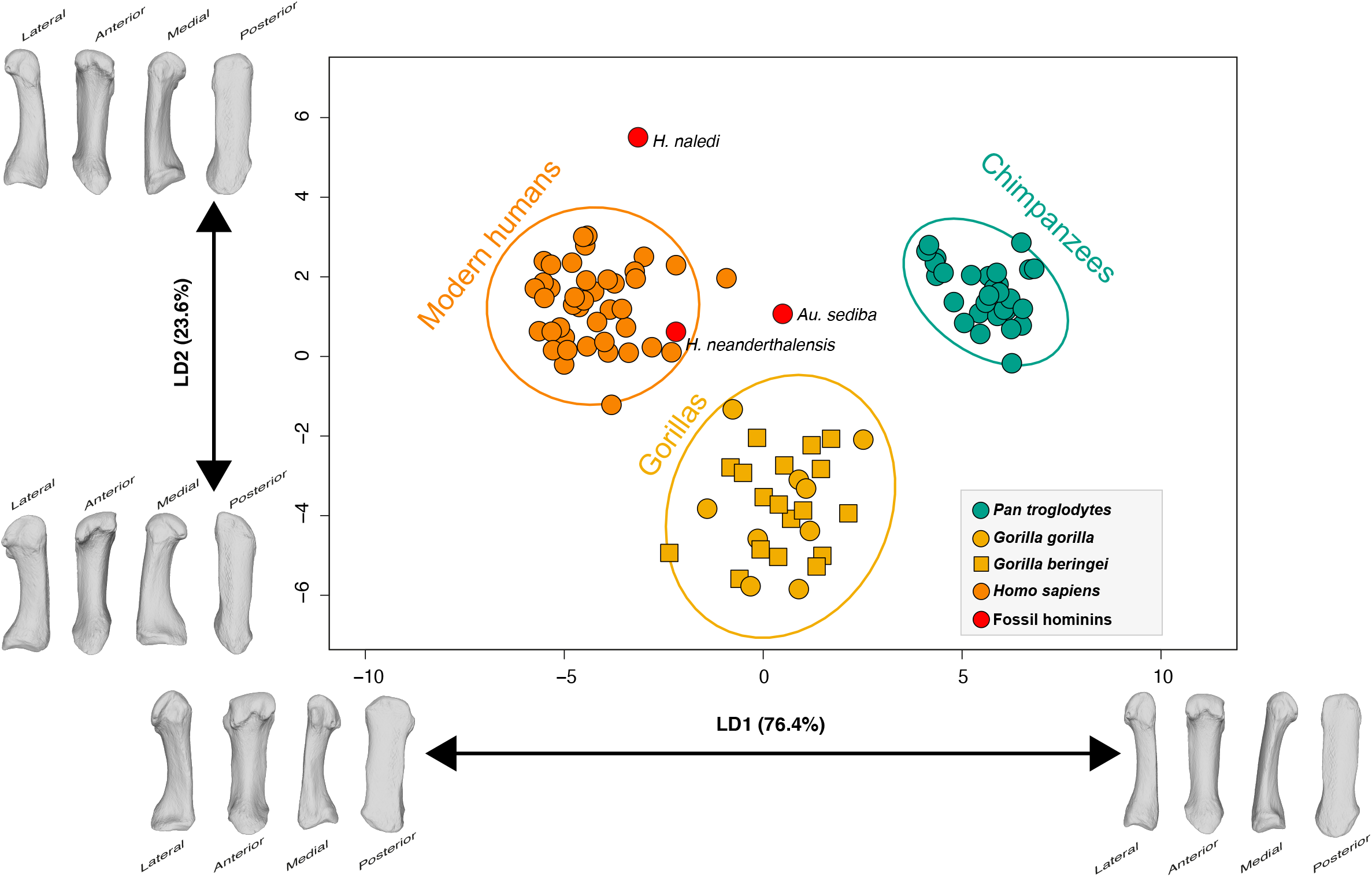
Multi-group linear discriminant analysis (LDA) of MC1’s shape using extant genera categories. One of the models closest to the mean shape was warped to match the multivariate mean using the thin plate spline method, then the obtained average model was warped to represent the variation along the two plotted CV axes.

### 3.5 Curvature

Table 5 provides the summed Menger curvature values for the six 3D models that were warped to represent the variation along the first three PC axes. These values were computed for both the dorsal (C1) and palmar (C2) sides of the MC1’s shaft and represent overall curvature. As expected, the palmar side of the shaft (C2) is more curved than its dorsal counterpart (C1). This means that the palmar curvature (C2) values are always higher as compared to the dorsal ones (C1) for all analyzed PCs. Overall, the shapes associated with the maximum values for each one of the three PCs corresponded to straighter shafts along the palmar side. In addition, the palmar curvature value (C2) for the minimum scores along PC1 correspond to the highest curvature (i.e., C2 curvature value for PC1 min; Table 5). The region of the morphospace that corresponds to this shape is occupied by the chimpanzees (Fig. 2b). In summary, the shapes associated with modern humans and gorillas are straighter, whereas the shapes that describe chimpanzees exhibit a more pronounced curvature along the palmar side.

## 4 Discussion

The first hypothesis was that the shape of the human MC1 would differ significantly from that of *Pan* and *Gorilla*, due to the variation in their manipulative capabilities and locomotive behaviors, and that *Gorilla* would show more morphological affinity with humans than chimpanzees. Overall, these analyses provide support for this hypothesis, confirming that there is indeed significant morphological variation between the extant great apes. We also found clear differences between chimpanzees and gorillas, with gorillas closer (i.e., more similar) to humans than to chimpanzees in PC1. This is due to their broader and more robust MC1s of gorillas and humans (i.e., broader shafts and expanded ulnar and radial epicondyles), as compared to the slender and more curved MC1s of chimpanzees (Table 5). The second hypothesis was that fossil hominin species *H. naledi* and *H. neanderthalesis* would exhibit an MC1 morphology more similar to humans than other great apes and *Au. sediba* an intermediate morphology between the African apes and humans. The results also support this hypothesis, as observed in the PCA and LDA plots (Figs. 2a and 3). However, it is important to note that even though the three fossils are more similar to the modern humans, they also display some distinct features, different from those which would typically be expected in extant *Homo*.

### 4.1 Hominin MC1 shape

The 3DGM data indicate that there is a distinctive suite of morphological traits that distinguish humans from chimpanzees and gorillas (Fig 3.). The main distinguishing traits are a straighter and more robust shaft (Fig. 2b and 3; Table 5) accompanied by larger radial and ulnar epicondyles and flatter distal and proximal ends of the body (Fig. 2c and 3). It is important to note that African apes are not a homogeneous group. For instance, gorillas show morphological affinities with humans relative to the shaft robusticity, although with a more proximo-distally curved shaft along its palmar side (Table 5), and more rounded distal-most and proximal-most portions of the MC1’s body. The PCA shows that chimpanzees are characterized by a slender and more gracile MC1, and this trait makes chimpanzees the most distinctive genus amongst extant taxa (Fig. 2b and 3). Chimpanzees are also characterized by an intermediate curvature of the radioulnar and dorsovolar ends of the shaft (Fig. 2c and 3). Interestingly, we found that chimpanzees display a more proximo-distally curved MC1 shaft compared with gorillas and humans (Fig. 3; Table 5). To our knowledge, this property has been only studied in phalanges 2-5, with those of chimpanzees being more curved than those of gorillas and humans (Stern, Jungers, & Susman, 1995; Susman, 1979). This curvature degree at the shaft has been usually interpreted as an adaptation for suspension and overall arboreal locomotion in digits 2-5 (Rein, 2011). As for the thumb, there is preliminary data that seems to indicate that it is routinely recruited during suspension in orangutans (MCclure Phillips, Vogel & Tocheri, 2012), yet its role has not been fully studied in chimpanzees. Consequently, it is not possible associate the MC1’s curvature observed in chimpanzees with the suspensory behaviors of this species.

As for the fossils studied here, results indicate that they all show a unique repertoire of morphological traits, different from those of extant genera. The general scientific consensus in recent years is that *H. neanderthalensis* had a hand morphology that was very similar to that of humans (Karakostis, Hotz, Scherf, Wahl, & Harvati, 2017; Karakostis et al., 2018; Niewoehner, 2001, 2006; Tocheri et al., 2008; Trinkaus & Villemeur, 1991). Our obtained results align well with this consensus, with the *H. neanderthalensis* specimen showing several similarities with the modern humans. The described morphology is one of a flatter (PC2) and broader (PC1) distal-most portion of the shaft, bigger epicondyles at the distal head (PC1) and a flatter proximal-most area of the shaft (PC2). However, *H. neanderthalensis* also differs in exhibiting a particularly robust MC1 with strongly marked muscular insertions, which distinguishes it from the rest of the sample, particularly along PC3. Neanderthals are known for the large flanges on their MC1s for the insertion of the opponens pollicis muscle that results in a waisted appearance of the MC1 in an anterior or posterior view (Maki & Trinkaus 2011; Trinkaus 1983). This trait also appears, to varying degrees, among some modern human populations, but rarely to the extent observed in *H. neanderthalensis* (Trinkaus et al. 2014). In our sample, this trait clearly distinguishes H. *neanderthalensis* from the rest of the analyzed specimens along PC3.

Previous reports indicate that the general morphology of the *H. naledi* MC1 aligns more closely with humans than apes, whilst still possessing a number of more primitive characteristics than the human MC1 (Kivell et al., 2015; Galletta et al. 2019). In our results we found that *H. naledi* aligns closer to humans in terms of shaft robusticity (Fig. 2b) and well-developed crest for the insertion of the opponens pollicis muscle (PC3) as it was reported by Kivell et al., (2015) and Galletta et al. (2019). However, it is also close to the range of morphological variability of chimpanzees in both PC1 and PC2, which indicates that the robusticity and curvature of the radioulnar and dorsovolar ends of the shaft is not similar to what is observed in modern humans. Even though the LDA robustly classifies *H. naledi* within the *Homo* category, it is worth mentioning that this specimen occupies a particularly unique position when projected to the LDA space. This is also observed in its position in the PCA, which seem to indicate an unusual morphology that can be described as displaying a narrower proximal end of the body, a relatively broader distal portion of the shaft, as well as marked attachment sites for the opponens pollicis and dorsal interossei. All these anatomical attributes contribute to generate the ‘pinched’ appearance of the palmar surface of *H. naledi’s* MC1 shaft (Kivell et al. 2015b).

Previous analysis of *A. sediba’s* hand morphology has found that it possessed several advanced *Homo*-like features, such as a longer thumb relative to shorter fingers, that potentially indicate advanced manipulative capabilities, whilst retaining primitive traits, such as a gracile MC1, similar to those of other australopiths (Kivell et al., 2011; Galletta et al., 2019). Our analysis showed that, unlike the Neanderthal and *H. naledi* specimens, *A. sediba* presented a general morphology that is more similar to chimpanzees than modern humans. *A. sediba* exhibits smaller epicondyles and a gracile shaft (Fig. 2b), with relatively flatter muscle attachments at the MC1 (Fig. 2d) than those observed in *Pan*.

### 4.2 Functional implications

Even though our study rigorously addresses the anatomical differences among the MC1s of extant hominines, any functional interpretations that we can advance are certainly inferred and not directly derived from our results. Hence, caution is required when interpreting these functional implications because shape differences could result from several different factors and not only be the result of different manipulative capabilities. Overall, our 3DGM results are consistent with previous assessments for the shaft morphology of the extant African apes and fossils hominins, and thus provides a morphometric support for the functional interpretations made based upon those features. The large epicondyles and robust shaft presented by the Neanderthal MC1 sample may suggest that they performed tool use in a very similar fashion to modern humans (Karakostis, et al., 2018; Niewoehner, 2001, 2006; Tocheri et al., 2008; Trinkaus & Villemeur, 1991). Nevertheless, the analyzed *H neanderthalensis* specimen also shows a classic Neanderthal feature (i.e., the opponens pollicis flange), which clearly distinguish it from the rest of the sample, particularly along PC3. It has been mentioned that it difficult to evaluate to what extent this trait may reflect muscle hypertrophy since the actual insertion area is mostly along the radiopalmar margin rather than across the palmar flange (Trinkaus, 2016). However, it is worth noticing that the radial extension of the opponens pollicis flange has been interpreted as increasing the opponens pollicis rotational moment arm, which suggests a greater mechanical effectiveness of this muscle in this species (Maki & Trinkaus 2011). *H. naledi* MC1’s anatomy suggests that this species was probably able to perform a certain degree of advanced manipulation, which might imply that this taxon was also a tool-user due to its robust shaft with marked muscular attachments but small epicondyles (Berger et al., 2015; Kivell et al., 2015a,b, Galletta et al., 2019). However, it is also worth considering that *H. naledi* shows an unusual MC1 morphology that when interpreted in combination with what is known from this species finger anatomy, may indicate a distinctive behavioral repertoire that could have included tool use as well as significant amounts of climbing (Kivell et al. 2015b). Finally, *A. sediba*’s anatomical characteristics suggests incipient tool using capabilities due to its slender thumb, smaller radial and ulnar epicondyles and curved joint surfaces (Kivell et al., 2011; Skinner et al., 2015; Galleta et al., 2019). Nevertheless, it is important to keep in mind that the above interpretations are exclusively based on a morphometric assessment of the MC1’s body anatomy (i.e., we did not directly assess any functional capabilities). Future studies should try not only to imply functional aspects based on morphological similarities but rather explicitly include them as part of the study (see e.g., Bucchi et al., 2020c).

From a functional perspective, the more robust MC1 shaft of humans (Fig. 2b) has been associated in previous studies with the ability of withstanding higher stresses placed upon the thumb by sustained power and precision grasping (e.g., Key & Dunmore, 2015; Marzke, Wullstein, & Viegas, 1992; Rolian, Lieberman, & Zermeno, 2011). These robust thumbs have also been related to a greater development of the thenar musculature attached to the shaft that is highly active during hard hammer percussion and favors thumb opposition (Marzke, 2013; Marzke, Toth, Schick, & Reece, 1998). The pronounced radial and ulnar epicondyles found at the distal head of the human MC1 (as described by PC1) may help to reduce the range of motion and stabilize the MCPJ (Imaeda, An, & Cooney, 1992). These epicondyles act as anchor points for collateral ligaments, which insert at the base of the proximal phalanx. Larger epicondyles are therefore thought to act as stronger anchors by providing a greater area for the collateral ligaments to attach to, helping to stabilize the MCPJ during the high forces that are experienced by the thumb during manipulation (Galletta et al., 2019). The flatter and larger distal articular surface in humans serves a similar purpose and has been interpreted as an adaptation that limits dorso-palmar motion whilst preventing radioulnar motion (Barmakian, 1992), thereby stabilizing the MC1 and facilitating forceful power and precision grasping.

### 4.3 Conclusion

The aim of this study was to quantify the morphology of the MC1 shaft in extant African hominoids, in order to better characterize its morphology. This characterization is not only relevant to better understand hominine anatomical differences and similarities, but also to provide further insights about its possible relationship with manipulative capabilities. This can facilitate more informed functional interpretation of fossil hominin morphology and contribute towards future studies linking morphology and function in hominin thumbs. Our study found that each taxon presented a unique repertoire of morphological traits, not present to the same extent in the others. Overall, the results obtained both aligned with and added to past functional interpretations of hominin morphology, thereby reinforcing the validity of 3DGM as a method of quantifying MC1 morphology and providing a deeper insight into the anatomy of the thumb in both extant hominids and fossil hominins. In addition, fossil MC1s are frequently fragmentary, and their epiphyses are often damaged. Hence, our applied approach which exclusively focused on the MC1 shaft might be particularly helpful in paleoanthropological contexts.

## Acknowledgments

We are grateful to the following curators and institutions: Emmanuel Gilissen (AfricaMuseum), Anneke H. van Heteren and Michael Hiermeier (Zoologische Staatssammlung München), Javier Quesada (Museu de Ciències Naturals de Barcelona), José Miguel Carretero (Universidad de Burgos). AB was partially funded by a Becas Chile scholarship, whilst TP was funded by the Leverhulme Trust Early Career Fellowship, ECF-2018-264. This study was funded by the research projects AGAUR 2017 SGR 1040 and MICINN-FEDER PGC2018-093925-B-C32. We are algo grateful to the two anonymous reviewers, associate editor and editorial board member who helped us to improve our work.

## Author contributions

Jonathan Morley: Data analysis; investigation; methodology; writing-original draft; writing-review and editing. Ana Bucchi: Conceptualization; methodology; resources; data curation; writing-review and editing. Carlos Lorenzo: Resources; data curation. Thomas A. Püschel: Conceptualization; data curation; data analysis; investigation; visualization; writing-review and editing; project administration.

## Data availability statement

The data supporting the findings of this study are available in the supplementary material (S6) of this article.

